# Amplitude Analysis of Polarization Modulation Data and 3D-Polarization Demodulation (3D-SPoD)

**DOI:** 10.1101/2020.03.10.986034

**Authors:** Andreas Albrecht, Dominik Pfennig, Julia Nowak, Rainer Matis, Matthias Schaks, Nour Hafi, Klemens Rottner, Peter Jomo Walla

## Abstract

Super-resolution optical fluctuation imaging (SOFI) is a technique that uses the amplitude of fluorescence correlation data for improved resolution of fluorescence images. Here, we explore if also the amplitude of superresolution by polarisation demodulation (SPoD) data can be used to gain additional information about the underlying structures. Highly organized experimental as well a simulated actin filament data demonstrate a principle information gain from this approach. In addition, we explored theoretically the benefits of analyzing the entire 3D-polarization information instead of only 2D-projections thereof. Due to fundamental principles, the probability of finding parallel orientations is approaching zero in 3D-SPoD in contrast to 2D-approaches. Using the modulation-amplitude based analysis we explored systematically simulated 3D-single molecules data (for which the true structures are known) under different conditions that are typically observed in experiments. We found that this approach can significantly improve the distinction, reconstruction and localization. In addition, these approaches are less sensitive to uncertainties in the knowledge about the true experimental point-spread-function (PSF) used for reconstruction compared to approaches using non-modulated data. Finally, they can effectively remove higher levels of non-modulated back-ground intensity.

## Introduction

In the past decades the detection of the polarization of light emitted by fluorescing labels as well as the polarization of the light used for excitation have proven to be very useful tools to obtain additional information about the structures to which the labels are attached ^1 2 3 4 5 6 7 8 9 10^. In addition, its use to disentangle subdiffractional structures has recently become the focus of several research groups (^11 12 13 14 15 16^). Several different ways for extracting different information from such data have been proposed (^11 12 13 14 15 16^) and it is not clear yet, what further information can be extracted from the heterogenous, homogenous or fluctuating orientation of the molecules that is intrinsically present in all fluorescence microscopy data. For example, using the additional orientation information for improvements in single molecule localization accuracy by considering the images from single molecules at out-of-focus position has only be explored very little so far, if at all, partly due to the vast computational effort that is necessary for such approaches. Obviously, still much of the molecular orientation parameter space that is intrinsically present in fluorescence microscopy data has not been explored yet.

Here, we explore the amplitude information in analyzed modulation data. We demonstrate experimentally that this can, in principle, provide additional information about subdiffractional structures in a similar fashion as in Super-resolution optical fluctuation imaging (SOFI).

In addition, we propose and explore theoretically 3-dimensional superresolution by polarisation demodulation (3D-SPoD). It is clear that one important limitation in polarization approaches is the inability to help with distinction between structures that exhibit nearly identical fluorescence polarization. However, using the full 3D-space instead of using only a 2D-projection can significantly reduce this limitation as the probability to find similar orientations in randomly oriented structures is much smaller in 3D.

This fundamental fact is illustrated in Figure 1 a-e. While Figure 1 a shows a random distribution of orientations, Figure 1 b shows only a subset of nearly parallel orientations in an angle range of *θ* = 0° ± 10° whereas Figure 1 c shows a subset of nearly perpendicular angles in a range *θ* = 90° ± 10° at an identical density as for Figure 1 b. It is immediately obvious that and why the probability to find structures of similar polarization in 3D is much smaller than to find structures that are perpendicular. The histograms showing the probability to find two structures spanning a certain angle in 2 or 3 dimensions in Figure 1 d and e are further illustrating this. While in a 2D-projection the probability to find two structures with parallel orientations (spanning an angle of *α*=0°) is the same as to find structures spanning any other angle (Figure 1 d), the probability to find two structures with parallel orientations is actually approaching zero in the case of a full 3D distribution but it is the highest for perpendicular angles (*α*=90°). In 3D less than 10-15 % of randomly oriented structures span a close to parallel angle even if a relatively broad range of 0°±30° is considered. In contrast, around 50 % span a nearly perpendicular angle of 90°±30°, at which structures can be differentiated very easily by polarization. Thus, it can be expected that any method that is based on discerning randomly oriented structures by their orientation yields more homogenous results throughout a sample when detecting the full spatial 3D orientation.

**Figure 1:**
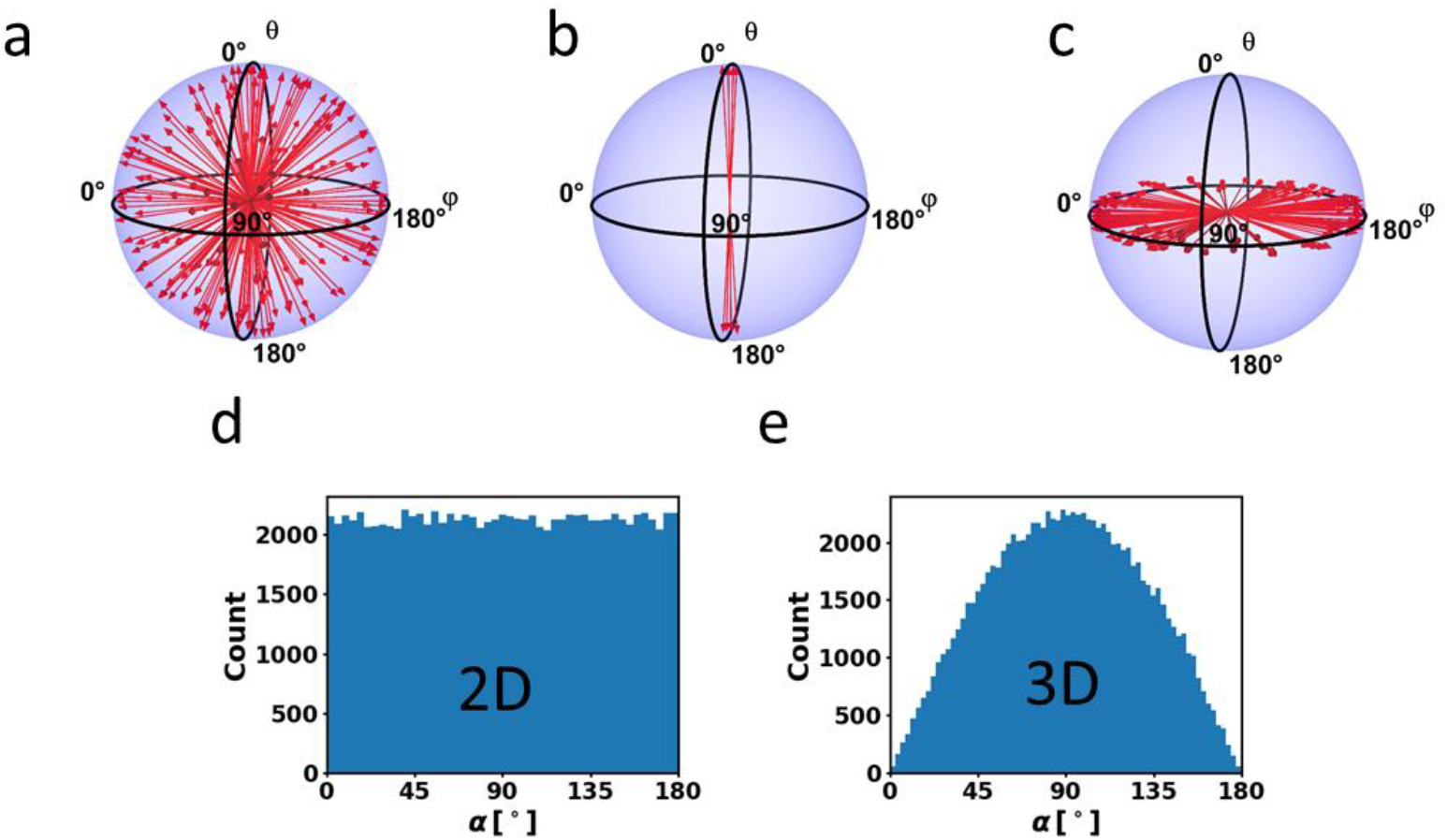
Representations of random orientations in space. a) Homogenous distribution of orientations, b) subset of a homogenous distribution of orientations with a polar angle less than 10°, c) subset of a homogenous distribution with a polar angle of (90 ± 10)°. For better visibility the arrow density in Figure 1 a is lower than in Figure 1 b and c . d,e) Histograms of angles *α* between 100000 pairs of randomly oriented vectors in 2D and 3D space respectively.

To detect the 3D-orientation we propose an experimental 3D-extension of our previous Super-resolution by Polarization Demodulation approach (3D-SPoD) and a method of analysis based on the modulation amplitude. We present simulated and experimental data that explore the ability of these approaches to disentangle and localize subdiffractional structures. These data demonstrate that they can significantly improve the distinction and localization, are more insensitive in uncertainties in the knowledge on the exact experimental PSF compared to deconvolution approaches using nonmodulated data and can effectively remove higher levels of non-modulated back-ground intensity.

## Results

### Modulation Amplitude based Analysis for experimental and simulated actin fibers

We first introduce the basics of the modulation amplitude analysis we propose for polarization demodulation data. For clarity, we will in the following refer to all data that shows diffraction limited raw data as *raw (image) data*, whereas data that describes the original underlying structure is called *original or underlying (image) data* and data that is obtained in any attempt to reconstruct the original images from raw images is referred to as *reconstructed (image) data*. Raw data can either be experimental or simulated data, where only in the latter case the original image is exactly known.

To analyze the data we propose a reconstruction that uses only the modulation amplitude components from the data. One feature of the analysis is based on the fact that any 2D-projection of polarization modulation signals as well as overlapping modulation signals in any pixel of all raw data, original or reconstructed image can all be fully described by the following function that contains three fitting parameters per pixel. These three parameters are the modulation amplitude, *A*, the modulation phase, *α*, as well as a constant offset, *I*_0_:

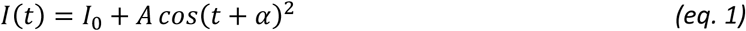

Let us briefly explore all scenarios of 2D-projections of experimental data that can be described by this function (Figure 2). The signal from a single emitter that is randomly oriented has a zero offset, *I*_0_=0, a certain amplitude, *A*, that depends on factors such as irradiation intensity and fluorescence quantum yield, and a phase-shift, *α*, that depends on the orientation of its transition dipole moment relative to the excitation polarization (Figure 2 a, b). If the emitter has a certain degree of rotational mobility, e.g. because it is not linked entirely rigidly to a labelled structure, this will lead to an increase in the constant offset, *I*_0_, at the cost of the modulation amplitude, *A* (Figure 2 c). Also, if several emitters are observed simultaneously in a distinct region of interest, their signal is described by eq. (1) because the sum of various cos^2^ functions with different phases, *α*, is again a cos^2^ function with lower amplitude, *A*, a higher offset, *I*_0_, and a single averaged value for the phase, *α*. If they have rather similar orientations, the reduction in *A* and increase in *I*_0_ will be smaller (Figure 2 d) than in a case of very different orientations (Figure 2 e). If unmodulated out-of-focus background is present this will only increase the constant offset, *I*_0_, but eq. 1 is still sufficient to entirely describe the signals (Figure 2 f). All these considerations are valid regardless of whether raw data or original unblurred data is to be described.

**Figure 2:**
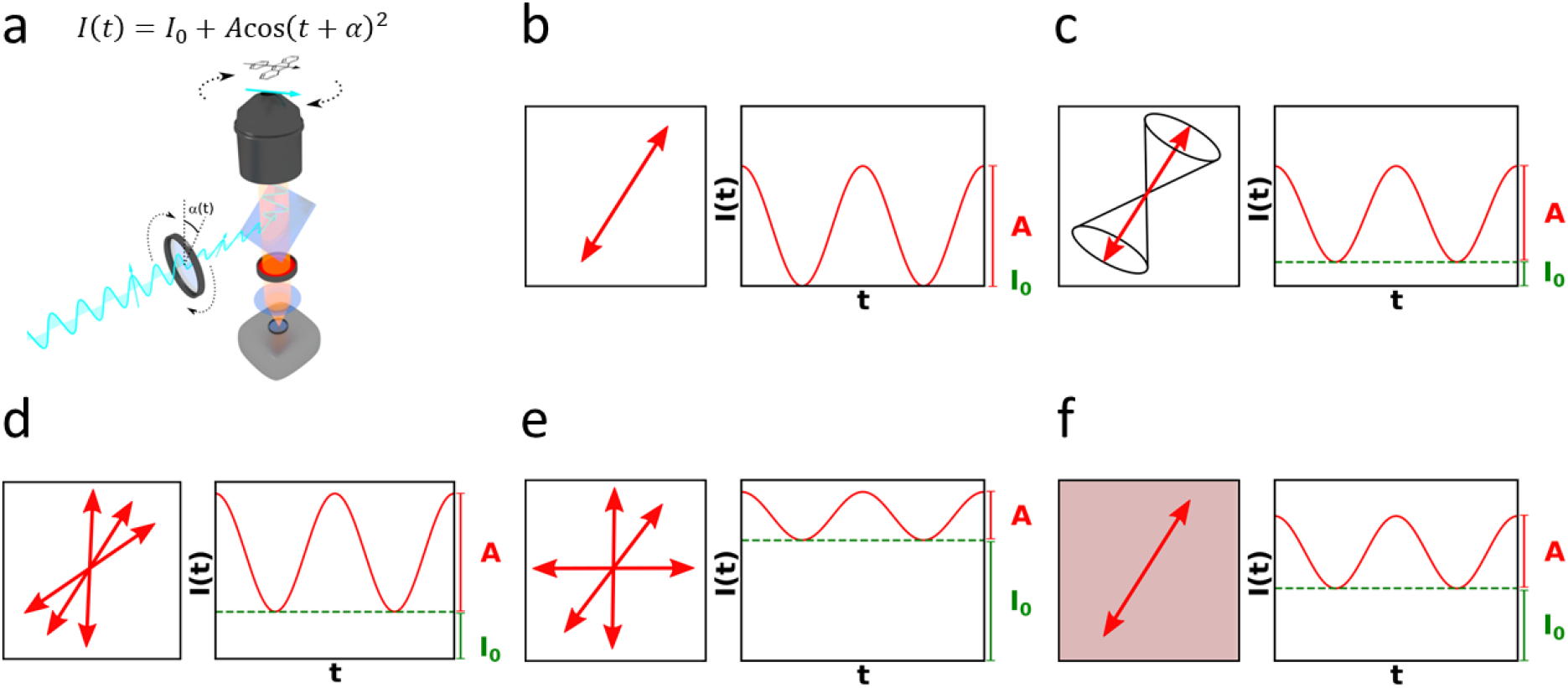
a) Schematic representation of the experimental setup used for data acquisition. b-f) Exemplary representations of transition dipole moment orientations and corresponding modulated fluorescence traces for b) a single, rigidly bound molecule, c) a single molecule exhibiting rotational mobility, d) a multitude of similarly oriented molecules, e) a multitude of differently oriented molecules, f) a single molecule with background fluorescence.

One advantage of using only the modulation amplitude for the analysis is based on is the fact that the oscillating fluorescence intensities exhibited by partly overlapping structures are phase-shifted to each other if the orientations of the labels’ transition dipole moments in the structure are different. Due to this phase-shift the modulation amplitudes in the overlap region are reduced (cf. Figure 2 d, e) resulting potentially even in a local minimum separating the structures in the amplitude image. Obviously this effect is more pronounced for bigger phase-shifts and accordingly bigger angles between the transition dipole moments. Another benefit of looking at the amplitudes is the intrinsic removal of background and noise that does not oscillate at the frequency imposed by the polarization modulation, this way much of the out-of-focus background is removed resulting in a sectioning capability similar to two-photon microscopy or SOFI.

Experimental raw data modulation movies are formed by blurring the signals emitted from the structures in each pixel of the original image – each characterized by the three parameters in eq. 1, modulation amplitude, *A*(*x*,*y*), the modulation phase, *α*(*x*,*y*), and constant offset, *I*_0_(*x*,*y*) – with the microscope’s point spread function, PSF. Here, *x* and *y* are the coordinates of the pixels (Figure 3).

**Figure 3:**
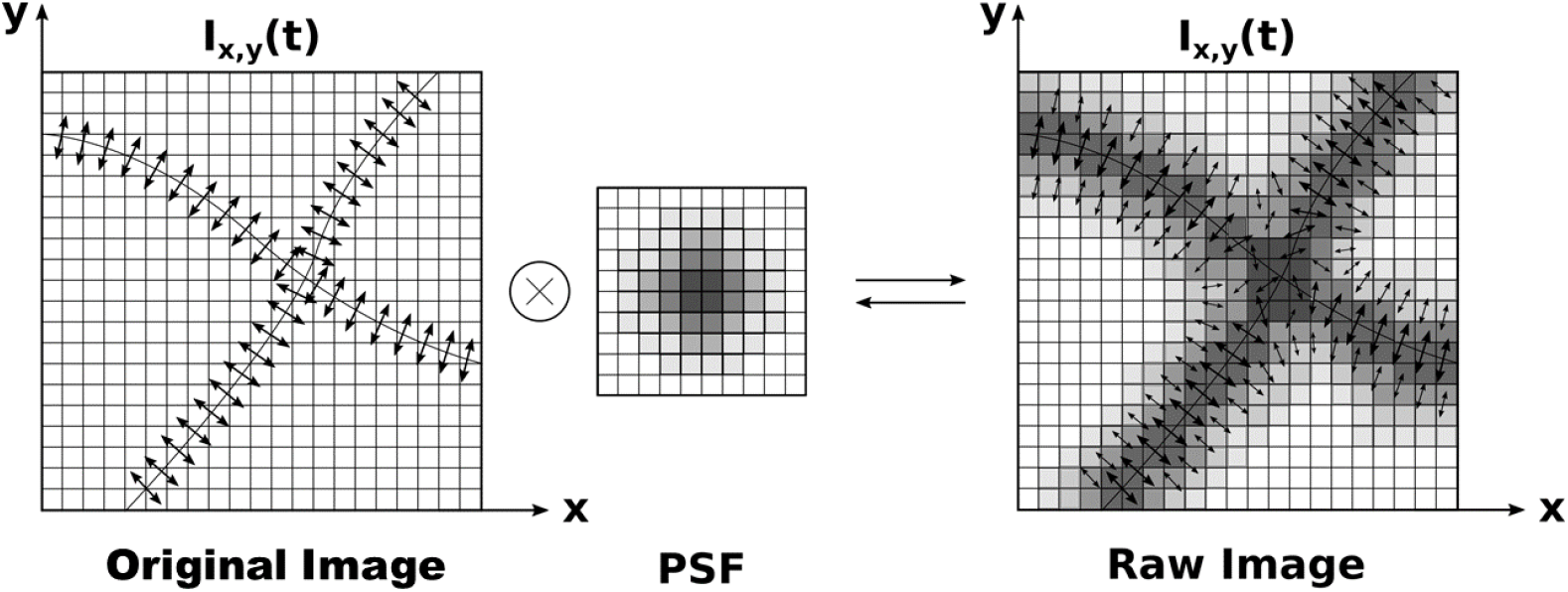
Schematic representation of relationship between original image and polarization structure, PSF and raw image and polarization structure.

As mentioned above, the reconstructed image can also be fully described by eq. 1’s three parameters for the signals emitted from the structures in each reconstructed pixel. The algorithm used here optimizes a corresponding parameter set (see methods section 2D-ALPA) in a way that if the images resulting from these parameters are blurred by the PSF, the sum of squared differences between this blurred reconstructed data and the measured raw data is minimized. All reconstructed data in this publication were obtained after 1000 iteration steps, unless noted otherwise. With this iteration depth no significant overdeconvolution could be observed in the analysis of the simulated data. The modulation amplitude based analysis to reconstruct the data requires no other parameter than the PSF and iteration depth. Currently the optimization is performed by using a nonlinear BFGS minimization algorithm^17^ and not by a faster linear optimization algorithm since eq. 1 introduces nonlinearities into the problem. Before each reconstruction a global, constant camera or image background noise level offset was subtracted from all raw data pixels. To test the amplitude approach, we explored experimental and simulated actin fibers in 2D-projections of the fluorescence polarization. In the following we will look at the reconstructed amplitudes *A*(*x,y*) and overall intensities *I*(*x*, *y*) = 0.5 . *A* (*x*, *y*) + *I*_0_ (*x*, *y*) (c.f Figure 2b-f).

Figure 4 shows raw data of the simulated and experimental actin fiber filaments at different relative angles. The simulated data were generated as described in the supplemental material wheras experimental data were taken from Reference ^18^. A common way to label actin filaments for fluorescence analysis, including live cell fluorescence microscopy, is the use of phalloidin conjugates^19 20^. Previous work has shown that when using rhodamine dyes to label actin filaments these dyes can have a very distinct orientation of the transition dipole moment relative to the filament (^18 21 22^). For example, when using Atto 590 phalloidin, apparently the transition dipole moments of all dyes are aligned perpendicular with respect to the orientation of the fibers resulting in modulation phases *α* that are directly correlated to the orientation of the fibers ^18^ (Supplementary Movie 1). Applying a color-coding to the image intensities that encodes the phases *α* in experimental and simulated data indicates that this allows the separation of pairs of fibers down to distances of about 50 nm even in raw data (Figure 4 e, i, o, r). Color-coding was done using a simple, custom ImageJ plug-in that allows choosing two different colors that are assigned to pixels with phases *α* above or below a selected value (Supplementary Software 1). In addition, intensities (Figure 4 f, j, p, s) and amplitudes (Figure 4 g, k, q, t) reconstructed with the analysis are shown. Selectively showing only phases in the reconstructed amplitude images that correspond either to one or another fiber (Figure 4 d and insets n) demonstrates that the individual fibers are reconstructed with high linearity, including the subdiffraction limited crossing region, for the simulated as well as experimental data. This is confirmed for reconstructed data that are color-coded as described above (Figure 4 h). A comparison of the reconstructed intensity image data and the corresponding pure amplitude data demonstrates the potential of the approach to use only the latter: The fibers are separated much better and more linear when using the pure modulation amplitude (Figure 4 g, k, q, t) than with the entire intensity (Figure 4 f, j, p, s). For differences in the polarization angle down to about Δ*α* = 38 ° and potentially below that the fibers can clearly be separated down to 50 nm (Figure 4 h). The color-coded presentation of raw data demonstrates that even fibers differing only by about 10° in angle can still be separated atdistances below 100 nm (Figure 4 u). However, it is important to note that this is facilitated here because often only two structures are to be separated. More areal structures with more than two overlapping features are considered in the 3D-single molecule simulations in the next section.

**Figure 4:**
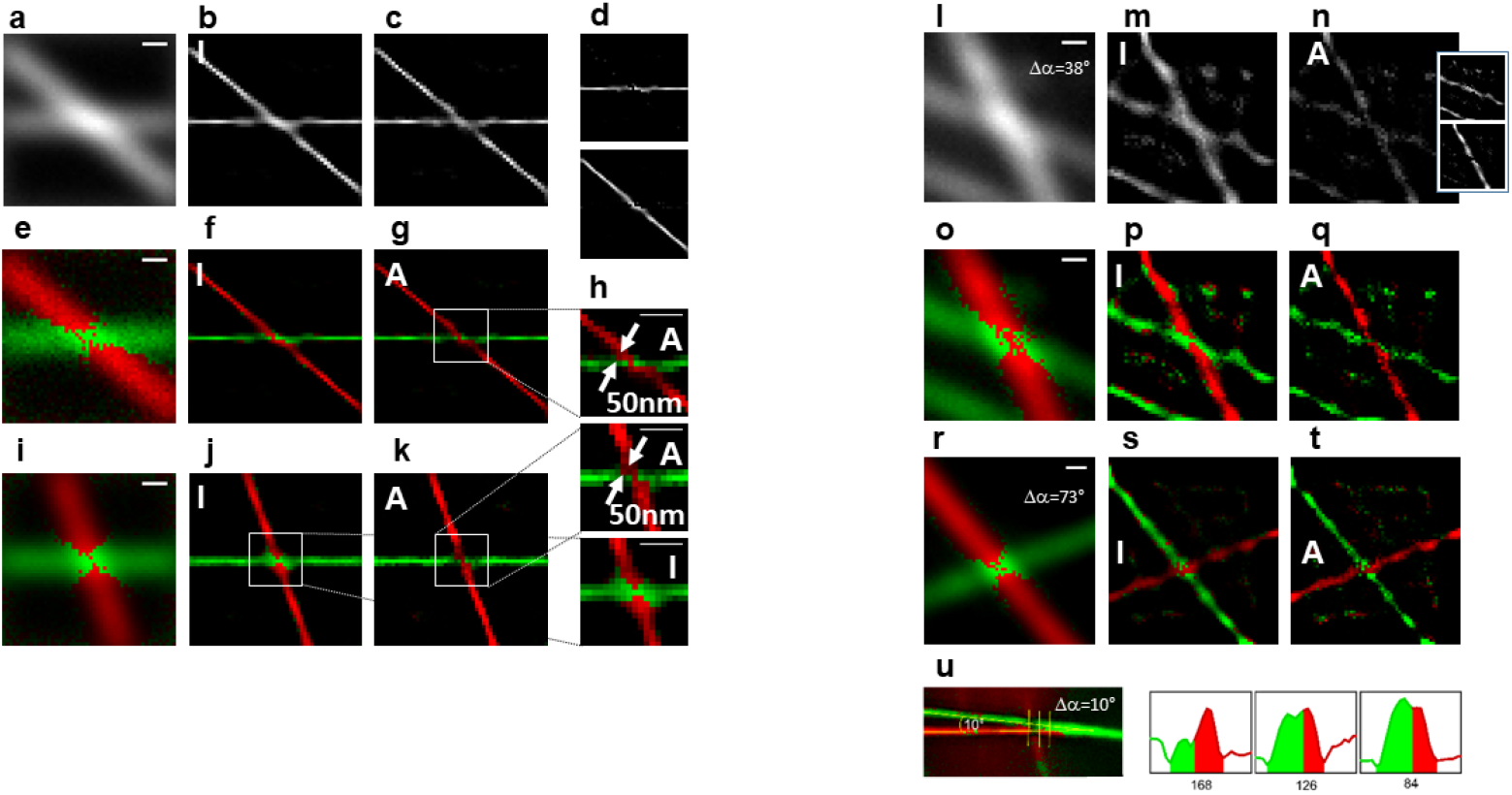
Pictures showing the analysis (here 2400 Iterations were used on the measured data l-u) of a-k) Simulated and l-u) measured actin fiber images. a,l) show the raw averaged images while e,o,i,r) show the colorcoded raw data. The reconstructed intensities are shown in b,f,j as well as m,p,s where f,j and p,s are two-color-coded. c,g,k) and n,q,t) show the reconstructed amplitudes where g,k and q,t are two-color-coded. d) and inset to n) singular isolated fibers. h) Magnifications of the crossing regions of the fibers with distance estimation.

### Analysis of the 3D-Polarisation and improvement in the homogeneity of the localization precision of single emitters distributed over a larger area

In order to detect the full 3D polarization the sample’s polarization must be observed under at least three different observation angles. The way that this can be achieved experimentally is by entering the back aperture of the microscope objective at three different positions, leading to three different light propagation vectors in the sample (Figure 5 a). To simulate areal structures with overlapping signals from several emitters we calculated a rectangular grid of 10 × 10 single molecules of random orientations but slightly subdiffraction-limited distances (Figure 5 b, Supplementary movie 2). This structure represents an areal square with a quite homogenous emission (Figure 5 b). Analysis of the 2D-SPoD signals observed for each of the three measurements corresponds to the projections of the transition dipole moments in the respective observation planes (Figure 5 a and c). As expected, there is heterogeneity in the reconstruction of each of the 2D projections. In addition, molecules illuminated from different sides are excited at different rates depending on their relative transition dipole orientations which can result in very different brightnesses seen in Figure 5 c. However, combining all three images by calculating the sum of the three 2D-amplitudes

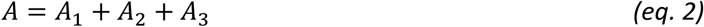

results in a significantly more homogenous representation (Figure 5 d).

**Figure 5:**
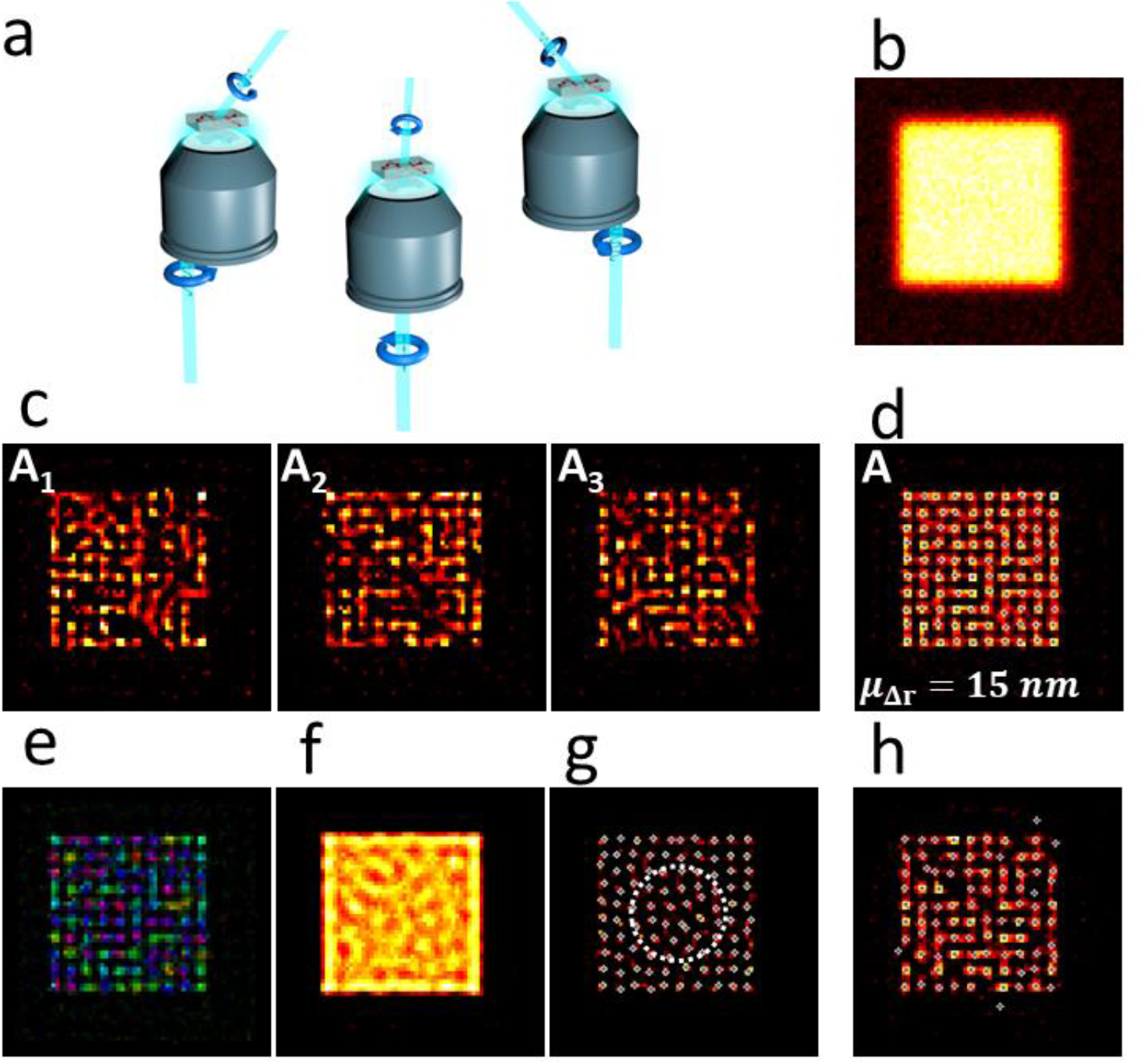
a) Schematic representation of the proposed illumination in 3D-SPoD showing the illumination beam entering the objective lens at three different positions resulting in three tilted illumination projections with rotating linear polarization. b) Average of the simulated 3D data with 100 emitters placed in a 10×10 grid pattern, c,d) 3D-amplitude analysis showing the three individual amplitude components A1,A2 and A3 (c) as well as the full 3D-analysis (d). The *μ*_*Δr*_ values show the mean value of the deviation *Δ*_*r*_ from the real simulated positions. The crosses indicate the positions where the emitters were localized. Details can be found in the methods section. e) amplitudes color-coded by phase f) Reconstruction using non-modulated data, g) Richardson-Lucy deconvolution of unmodulated data, h) Analysis of only one 2D-projection.

We quantified the homogeneity by fitting a manifold of 2D Gaussians to the analyzed data. The fitting is performed by using a custom localization script. The localizations are then compared to the real positions at which the simulated molecules are supposed to be found. The geometrical distance between real position and localization is averaged (*μ*_*Δr*_ in Figure 5 d, details in Methods section). While resolution depends on many factors such as measuring times, emitter intensities, orientational flexibility, knowledge of the experimental PSF and out-of-focus background, in case of the parameters given for figure 5 d (7500 photons per emitter) an average localization precision equivalanet to approximately ±15 nm is observed. Figure 5 e is a color-coded representation of Figure 5 d showing the different orientations (details in methods section) of the molecules.

Using non-modulated data in the 3D-ALPA analysis (Figure 5 f) or in a Richardson-Lucy-Deconvolution (Figure 5 g, Supplementary Movie 2 c, d) or only 2D modulated data for the ALPA analysis (Figure 5 h) yields significantly worse results. If using non-modulated data, the 3D-ALPA analysis provides no separation of the single emitters (Figure 5 f). Using non-modulated data in a Richardson-Lucy-Deconvolution (Figure 5 g, Supplementary Movie 2 c, d) leads to fragmentation that represents the real structure only partly. Even though the Richardson-Lucy-Deconvolution yields more or less acceptable molecule positions at the edges of the structure (Figure 5 g), in the more central part (highlighted by white, dashed circle) the estimated molecular positions do not represent the underlying structure. Finally, using only 2D modulated data for the ALPA analysis also leads to a smaller number of subbdiffractional details resolved (Figure 5 h). Please note that identical average photon counts per molecule were simulated whenever direct comparisons are shown throughout this entire work. This includes the direct comparison of Figure 5 f, g, h and 5 d. Of course, eq. 2 implies that the average per molecule countrates of A1, A2 and A3 in Figure 5 c are only a third of the sum shown in Figure 5 d.

It is obvious that when 3D-modulated data is used the molecules are localized more accurately than after a standard deconvolution. These comparisons were done with the assumption that the real PSF is exactly known. However, in reality it is nearly impossible to know the exact PSF for a given experimental set-up which is why most deconvolution based approaches remain very limited for real resolution enhancement.

### Results with uncertainties in the known PSF

As mentioned in the previous section, in most cases the real PSF is not exactly known for a given experimental set-up. If the PSF were known exactly, most deconvolution approaches would yield very reasonable results. However, the precise PSF is rarely known. Therefore, we also explored how sensitive the results are for uncertainties in the PSF and if the 3D-SPoD modulation approach can help in the reconstruction of the original structure without precise knowledge of the PSF.

Figures 6 g and h confirm that even a FWHM-deviation of only 10 % in an estimated PSF from the real PSF has a dramatic impact on deconvolution based reconstructions using non-modulated data. In contrast to Figure 5 g, Richardson Lucy deconvolution yields no single emitter reconstruction with such a PSF even after 5000 iterations (Figure 6 g). Increasing the iteration depth to 100.000 seems to reconstruct single emitters at first glance. However, the localization precision *μ*_*Δr*_ is larger than ±125 nm, which is larger than half the distance between the simulated structures(Figure 6 h). On top of that, in contrast to the Richardson-Lucy reconstruction of unmodulated data with correct PSF (Figure 5 g) it can happen that it resolves only 9 apparent molecules at the edges of the structure instead of the true number of 10 molecules.

**Figure 6:**
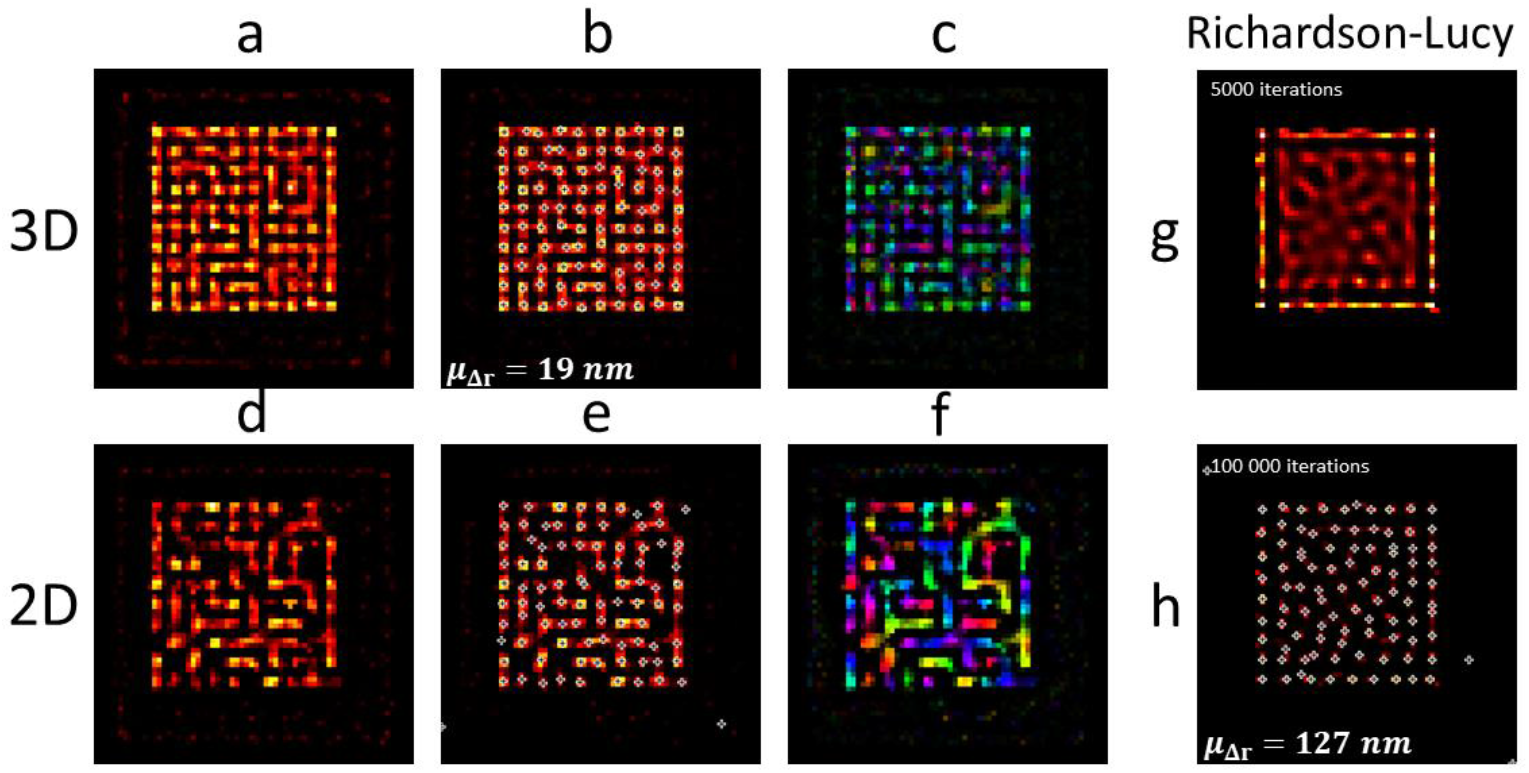
The a,d) reconstructed intensities b,e) amplitudes and c,f) color-coded amplitudes for amplitude reconstruction from 2D and 3D modulated data with an incorrect PSF (FWHM of 2D-Gaussian 10% too large), g, h)Corresponding Richardson-Lucy deconvolution of unmodulated simulations after 5000 (g) and 100000 (h) iterations. Again the localizations (crosses) and a mean value of the mislocalizations are shown.

However, when 3D-SPoD data are reconstructed using 3D-ALPA with the same perturbed PSF, the underlying structure can still be recovered reasonably well (cf. Figure 6 a, b). The following localization precision is with ±19 nm only slightly worse compared to the reconstruction with exact known PSF (Figure 5 d). This result demonstrates the potential of the 3D-modulation approach to use the different phases of different emitters for disentanglement and correct localization even under experimentally conditions with uncertainties in the known PSF. When using only 2D-projections of modulated SPoD data, again only a smaller number of subdiffractional details is reconstructed (Figure 6 d, e) since fewer features can be excited at significant rates.

### Sectioning capability and orientational flexibility

In real imaging applications, e.g. cell imaging, usually significant out-of-focus fluorescence is also present that severely hinders detailed analysis of the fine structure in the focal plane. However, similar to SOFI^23^, the modulation amplitude decreases very quickly outside the focal plane as the number of structures contributing to overlapping out-of-focus signals increases rapidly. In other words, the out-of-focus background is rather unmodulated and contributes mainly to the constant offset *I*_0_ in the analysis (Figure 2 f) but not to the amplitude output image of the reconstructed data. To explore the influence of an irregular out-of-focus background we repeated the simulation and analysis of the 10×10 molecular Grid shown in Figure 5, adding a background gradient with a maximum intensity about 10 times higher than the single molecule intensities themselves (Figure 7 a). Supplementary movie 3 demonstrates that the signals seem to be severely impacted by such a background which is not untypical in fluorescence microscopy. However, applying the exact same analysis as described previously to such data demonstrates that using 3D-SPoD a reconstruction of the original structure with almost the same localization precision (±22 nm) as in the absence of such back ground is still possible (Figure 7 d). An analysis using unmodulated data is neither capable of removing unmodulated background nor of resolving the individual emitters (Figure 7 b and Supplementary Movie 3 c, showing the results after 1000 and 100000 iterations, respectively). This is due to frequency-modulated signal filtering similar to lock-in amplification. Signals with a constant background of 90% relative intensity represent also situations in which the modulation depth is greatly reduced due to orientational flexibility of fluorescence labels (compare Figure 2 c and f). Also in such situations different phases still distinguish different emitters, even if they are quite orientationally flexible.

**Figure 7:**
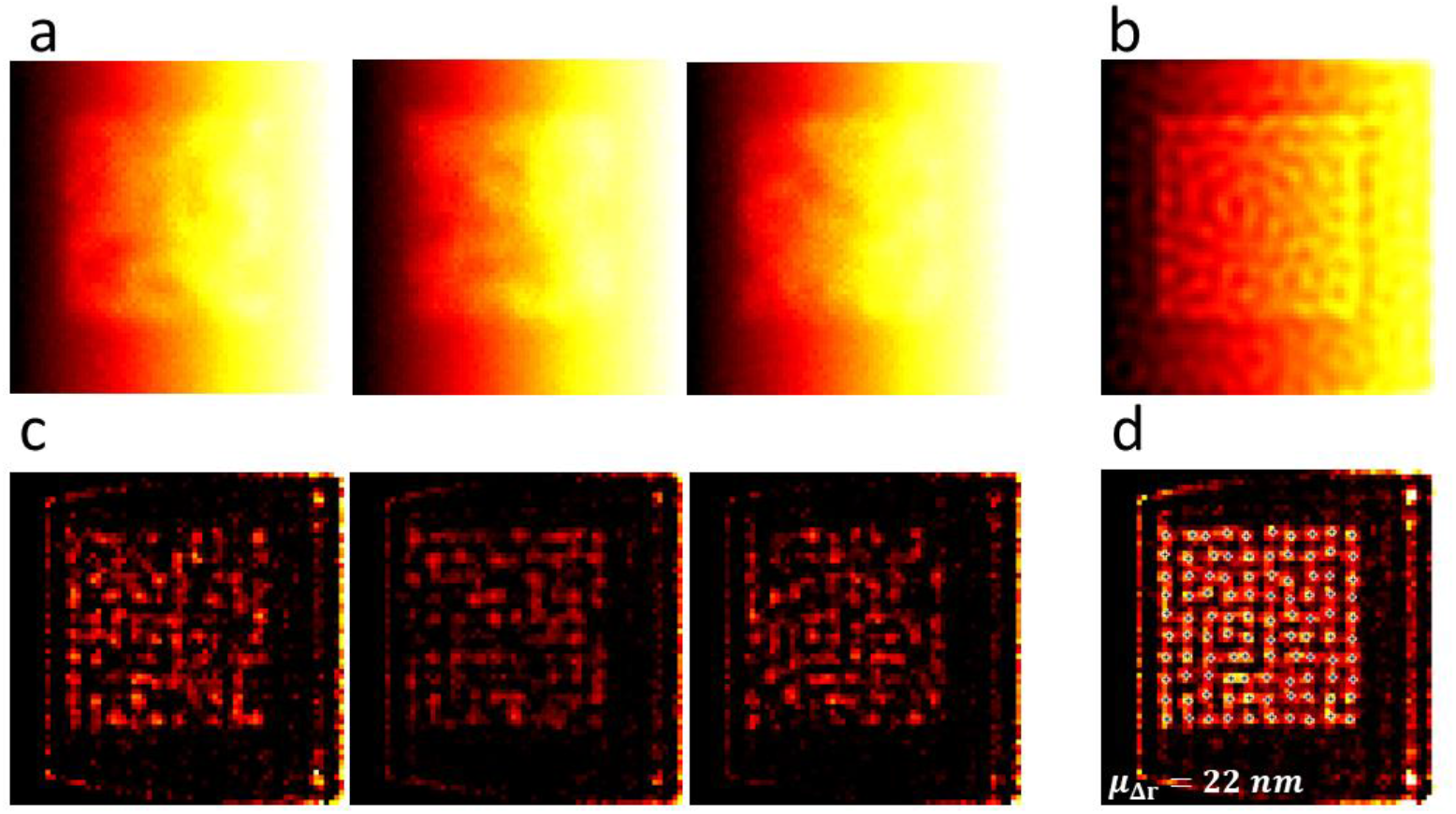
a) Temporal average of raw movies simulated for three illumination directions with a background linearly increasing from the left to the right. In the maximum the background is ten times more intensive than a single emitter, b) Richardson-Lucy deconvolution (1000 iterations) of the sum of the raw data, c) modulation amplitudes of the 3D data for each illumination direction after reconstruction, d) sum of the single reconstructed amplitudes. Again the localizations (crosses) and a mean value of the mislocalizations are shown. See also: Supplementary Movie 4 c.

## Summary

In the present report we have demonstrated that using only the modulation amplitude of analyzed polarization modulation data can, in principle, provide valuable additional information about subdiffractional structures (Figure 2 and 4). In addition we demonstrate that using polarization modulation in three dimensions (3D-SPoD) can significantly improve the distinction and localization of such structures (Figure 1 and Figure 5). While in a 2D-projection the probability to find two structures with parallel orientations (spanning an angle of *α*=0°) is the same as to find structures spanning any other angle (Figure 1 d), the probability to find two structures with parallel orientations is actually approaching zero in the case of a full 3D distribution (Figure 1 e). We explored simulated single molecule data under various conditions similar to the conditions typically observed in single molecule localization experiments. This allowed to compare quantitatively the deviation of the analyzed position from that of the true positions under such conditions. We propose a three-dimensional extension of SPoD set-ups (Figure 5 a) and an analysis that uses only the three modulation amplitude components of such 3D-SPoD data (3D-Analysis of local Polarization Amplitude, 3D-ALPA). Experimental and simulated data of actin filaments demonstrates that the amplitude based analysis can provide good linearity in reconstructed images of fiber crossings at subdiffractional distances (Figure 4). Simulations of rectangular areal structures represented by grids of 100 single molecules (7500 photons total per molecule) with slightly subdiffractional overlapping emission demonstrate that 3D-SPoD can localize all single emitters in parallel and with good precision (Figure 5 d) whereas unmodulated data did not allow separation of these subdiffractional details even if the exact PSF were known (Figure 5 f and g). If an experimental PSF were only known with a precision deviating by 10 % from the real PSF the capability to resolve or localize the molecules using unmodulated data was even worse (Figure 6 g,h) while 3D-SPoD data still allowed to resolve and localize the emitters (Figure 6 b) under otherwise identical conditions. In addition, simulating the presence of up to 10 times more intense out-of-focus background demonstrates the principle sectioning potential of SPoD and the ability to still detect and localize the single molecules with a similar precision (Figure 7d and supplementary Movie 3).

Even though significant future work is still necessary to explore the full molecular orientation parameter space that is intrinsically present in fluorescence microscopy data our results show that the approaches explored here can significantly improve separation and localization, provide reconstruction that is more insensitive to uncertainties in the PSF than deconvolution with unmodulated data and remove high levels of constant fluorescence background.

## Methods

### Two-dimensional polarization modulation measurements

The two-dimensional measurements were performed as previously described by Hafi et al. ^11^. The atto 590 labeled actin fibers are illuminated using a modelocked Ti:Sa laser (Chameleon Ultra II, 80 MHz, Coherent) and an optical parametric oscillator (OPO, APE) transforming the 800 nm laser output to 590 nm light. The beam is expanded using a telescope system. Afterwards the linearly polarized light passes a constantly rotating 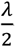-waveplate that rotates the polarization plane. The plate is rotated using a chopper system (Optical Chopper System, Thorlabs) that is synchronized to the EMCCD camera used for detection. A lens is used to focus the beam on the back aperture of the objective lens (NA = 1.35 oil immersion, UPlanSApo, 60x Olympus). The objective lens is mounted in an inverted microscope body (IX 71, Olympus). A dichroic mirror (dual line beamsplitter zt 488/594 RDC, AHF Analysentechnik, Tuebingen, Germany) is used to separate the emission from the excitation. After passing another telescope the light is detected by the EMCCD camera (iXonEM+897 back illuminated, Andor Technology).

### Analysis by local Polarization Amplitude (ALPA) Algorithm

The algorithm used for evaluation uses the model described in eq. 1 to describe the fluctuating intensity *I*(*t*) in each pixel. For better convergence of the algorithm eq. 1 has been reformulated to:

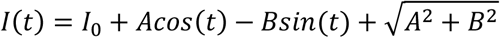

where the square-root term shifts the trigonometric part of the function towards non-negative values. The offset values *I*_0_ are set to zero if they would take negative values in any iteration. The individual pixels described by this three-parameter approach are then spatially blurred by the PSF to generate the recovered blurred data that is then compared to the measured raw data in a least-squares functional. This functional is then minimized by the BFGS algorithm from Python’s scipy library^17^. The starting parameters are determined from the Fourier transforms (also from the scipy library) of the raw data in each pixel. Similarly the amplitudes presented in this paper were generated by Fourier transforms.

We note that we could observe small differences in the obtained results when running the algorithm on Linux and Windows machines probably due to numerical noise. An alternative to Figure 4 is shown in Supplementary Figure 1 where the former was created on Windows 7 and the latter on Ubuntu 14.04.5. All other results were obtained using the Ubuntu machine.

### Three dimensional single-molecule orientation measurements with three-directional illumination

When the sample is illuminated from three sides the orientation can be recovered by considering the three planes that are being spanned by the respective light’s propagation vectors and the polarization vectors at fluorescence maximum in each pixel or for each emitter.

The orientation of the transition dipole moment can now be found by finding a vector that lies within all three planes. This vector can be found by finding a vector perpendicular to all three normal vectors of the planes which can be achieved by solving a system of linear equations where the dot products of the normal vectors and the unknown orientation vector are set to zero. The system was solved by using the “lsq_linear” function from the scipy library^17^.

#### Color-coding

To apply the color-coding to the images the modulating videos were Fourier transformed in each pixel along the temporal axis. An ImageJ plugin then assigns a color to each image pixel depending on the phase value found in the Fourier transformed image. In the two-color-coding the assigned color is a specified one (here: red or green) where a threshold value for the phase determines which color is assigned in each pixel (Figure 4). In the multi-color-coding the color is determined by proportionally mapping the phase values onto the “hue” value of the HSB color definition (Figure 10).

To visualize three-dimensional orientations the color-coding was applied analogously using the following value Φ(x, y) to define the color:

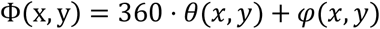

where *θ*(*x*, *y*) and *φ*(*x*, *y*) are the polar and azimuthal angles calculated for each pixel.

#### Simulations

The simulations were created on the basis of eq. 1. In the case of the fibers the phase was set to specified values corresponding to the fiber orientation, while the three phases of the emitters in the 10 by 10 grid were calculated from randomly generated orientations. The single emitters were blurred by a simple symmetrical 2D Gaussian function. Shot-noise was applied to the simulated photons (7500 total per single molecule, all molecules were simulated at identical total brightness). Additionally, normally distributed noise was applied to account for detector noise. The standard deviation was chosen to match the one found in measured data. The values for brightness and noise-level were chosen to correspond to values typically observed in a measurement of 1.8 s. Anisotropic emission by the molecules has not been taken into account. This would cause molecules that lie flat on a surface to appear brighter than those perpendicular to the surface.

#### Localization and Homogeneity estimation

In order to quantify the homogeneity of the resolution in the simulated data the ALPA-analyzed data was compared to the known underlying structure. In order to localize the simulated emitters the localization algorithm first looks for the brightest pixel in a range of the FWHM of the PSF around the true molecule positions. The 100 pixels determined in this way are then used as starting positions for a fit of a sum of 100 two-dimensional Gaussian functions. The fit is performed by scipy’s “least_squares” function ^17^. The distance between the true position and the calculated position is determined for each emitter. Finally, all 100 distances are averaged to yield *μ*_*Δr*_.

#### Preparation of linear actin filaments

The data for the linear actin filaments was taken from ^18^ so the preparation procedure is identical [as described therein]: 250 *μg* of rabbit skeletal muscle G-actin (AKL99-A; Cytoskeleton, Inc., Denver, CO) were resuspended to 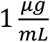 using 250 *μL* of General Actin Buffer (BSA01; Cytoskeleton, Inc., Denver, CO) containing 0.2 *mM* ATP (New England Biolabs Inc., Ipswitch, MA) as well as 1.0 *mM* DTT (Carl Roth, Karlsruhe, Germany). After stirring the mixture was cooled on ice for 1h. 5 *μL* of 10x polymerization buffer solution (BSA02; Cytoskeleton, Inc., Denver, CO) mixed with 45 *μL* of the previously produced solution were left for 1h at room temperature to polymerize. Afterwards the solution was diluted 100-fold using 1x polymerization buffer containing 70 *nM* Atto590-phalloidin (ATTO-TEC, Siegen, Germany). A microscope coverslip was spotted with 10 *μL* of the solution containing the actin filaments. After the solution spread out, additionally 15 *μL* of ProLong Gold Antifade Mountant (Life Technologies, Darmstadt, Germany) were spotted in the center on the coverslip. Before measurement the sample was left alone for more than 30 minutes.

## Acknowledgements

This work was financially supported by the Deutsche Forschungsgemeinschaft (DFG) (INST 188/334-1 FUGG) and the DFG-funded Research Training Group „Protein Complex Assembly“(GRK2223/1).

