## Supplemental data for "Amplitude Analysis of Polarization Modulation Data and 3D-Polarization Demodulation (3D-SPoD)"

### Slide 1
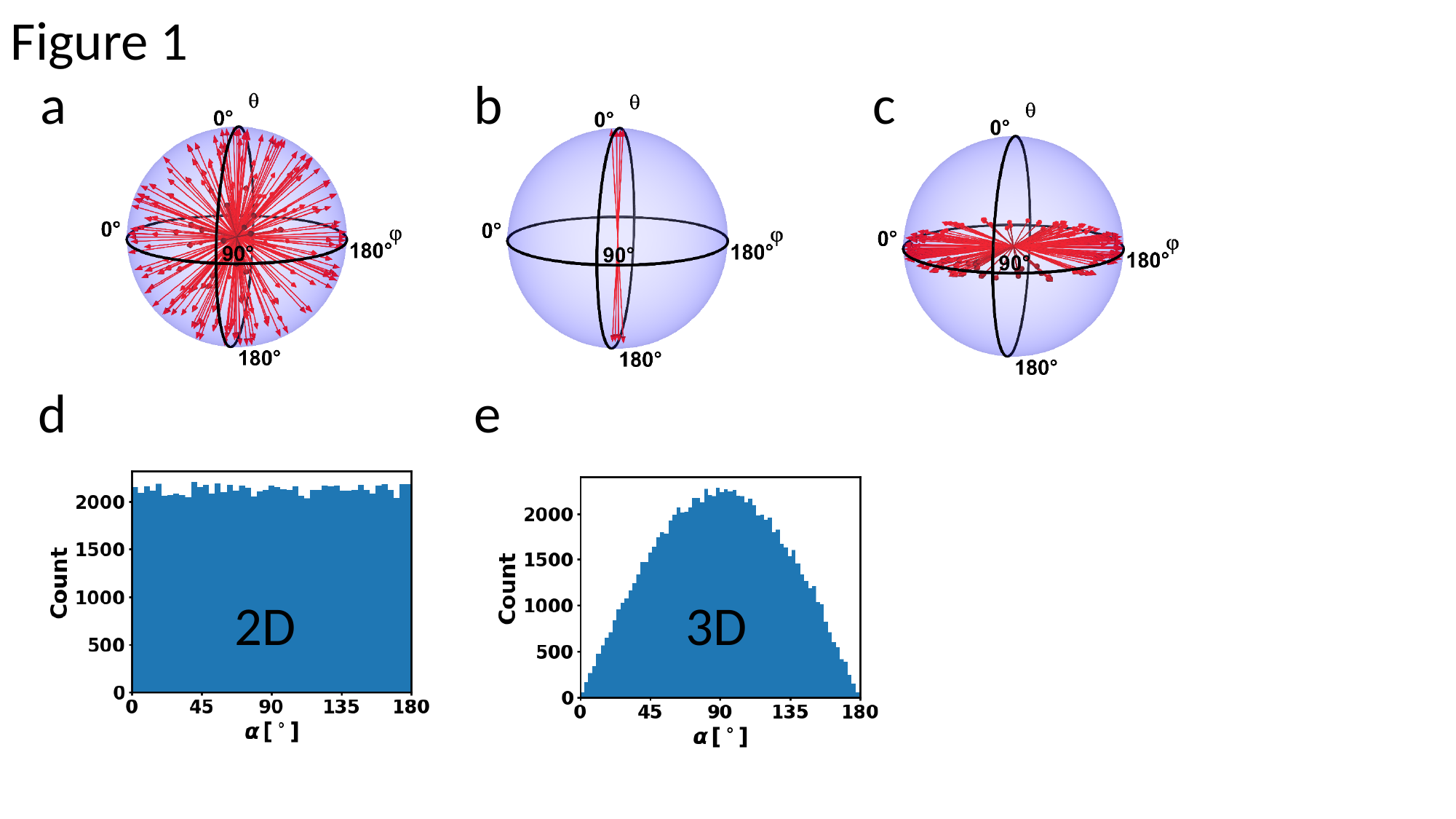

Figure 1
a b c
d e
2D
3D

### Slide 2
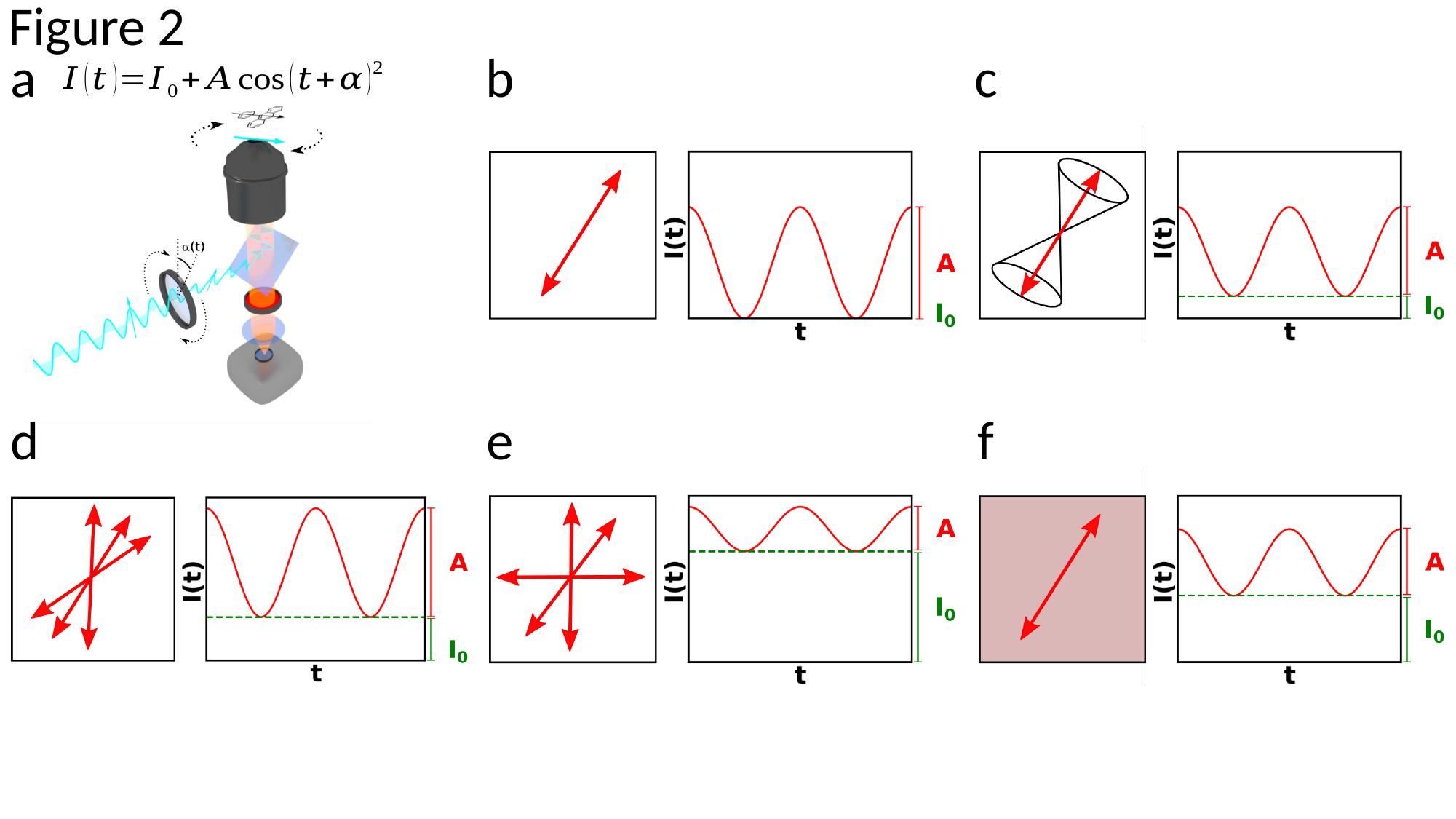

Figure 2
b
c
a
e
f
d

### Slide 3
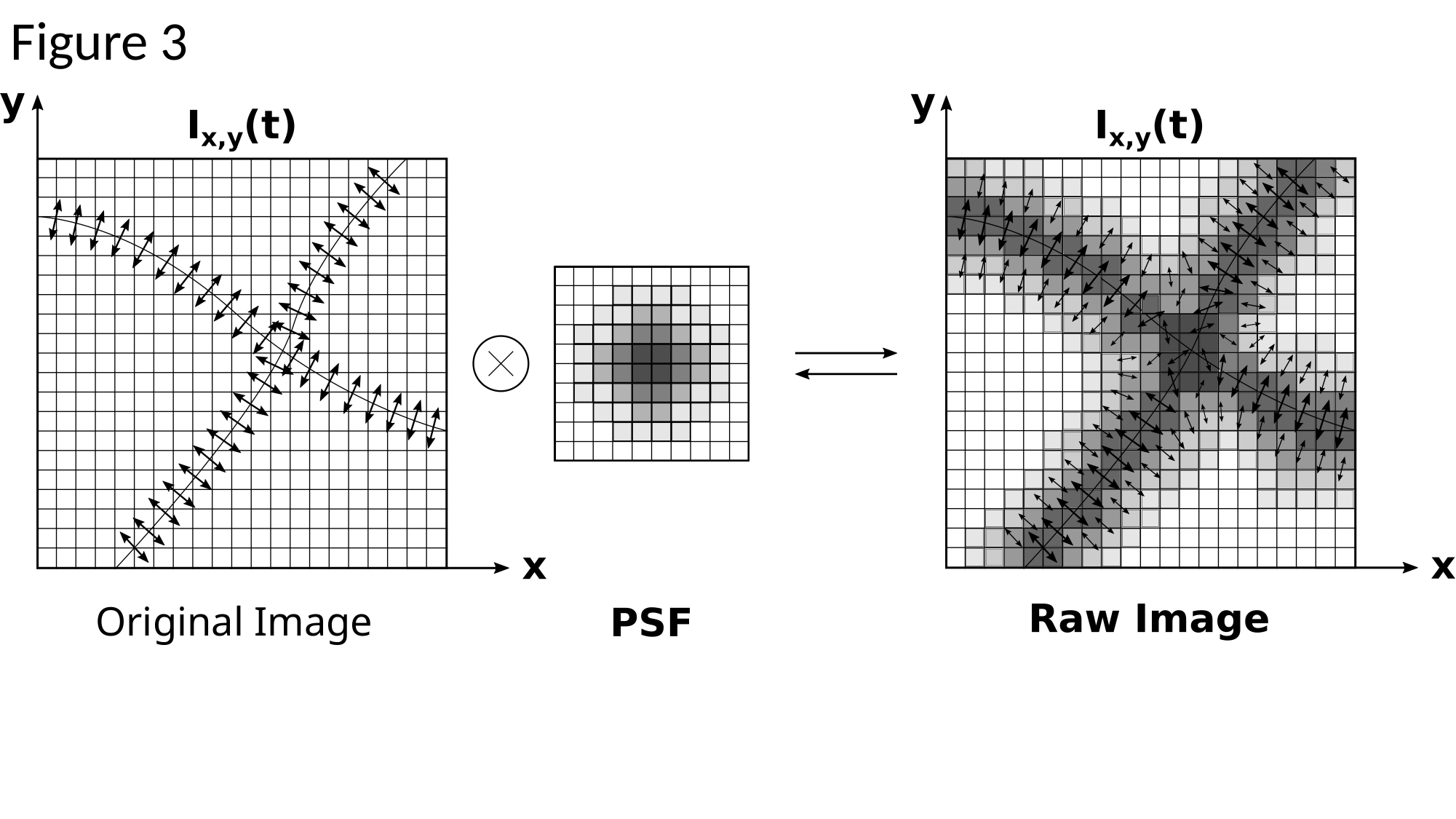

Figure 3
Original Image

### Slide 4
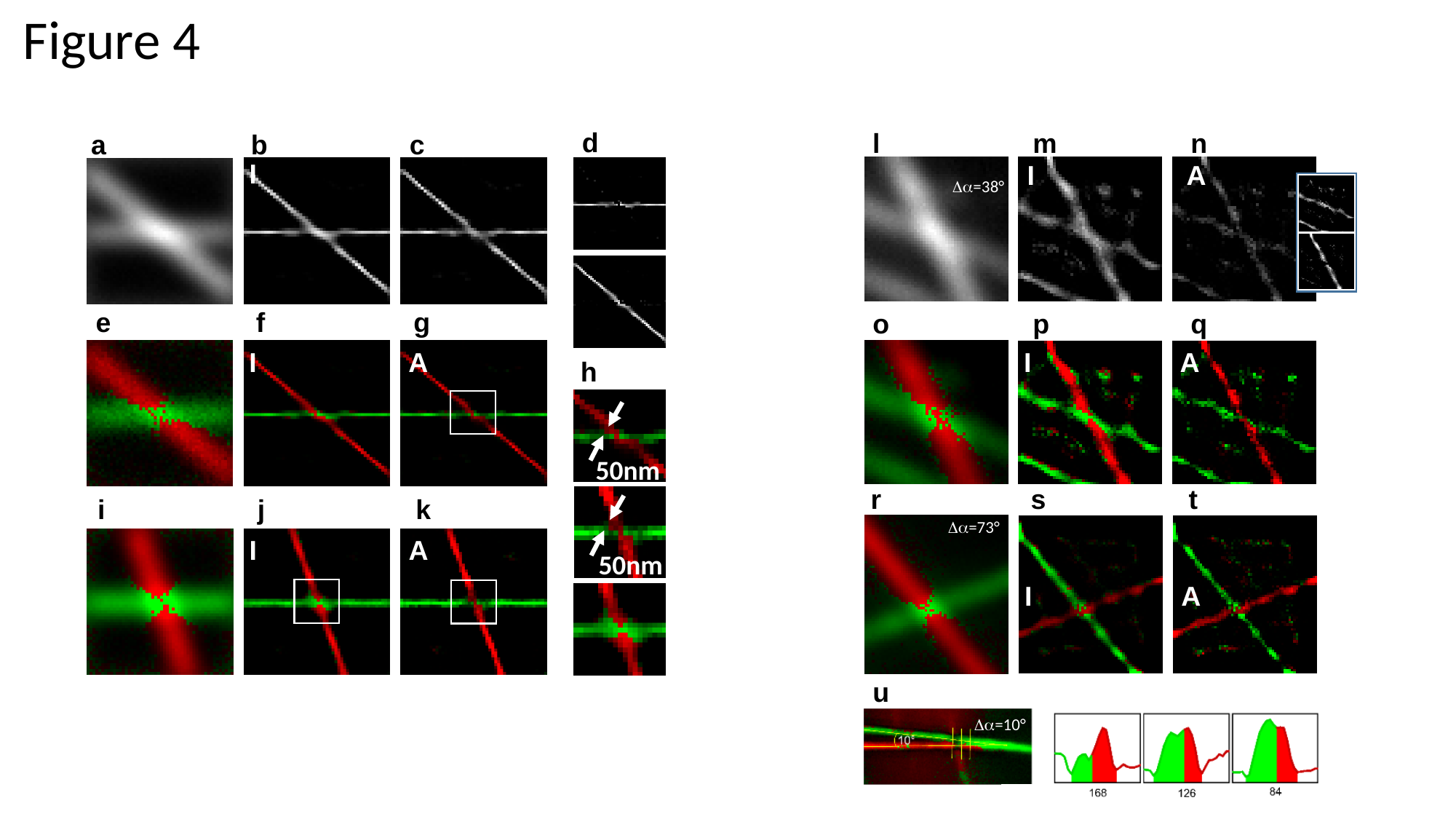

Figure 4
d
l
m
n
a
b
c
I
I
A
Δα=38°
e
f
g
o
p
q
I
A
I
A
h
50nm
r
s
t
i
j
k
Δα=73°
I
A
50nm
I
A
Δα=10°
u

### Slide 5
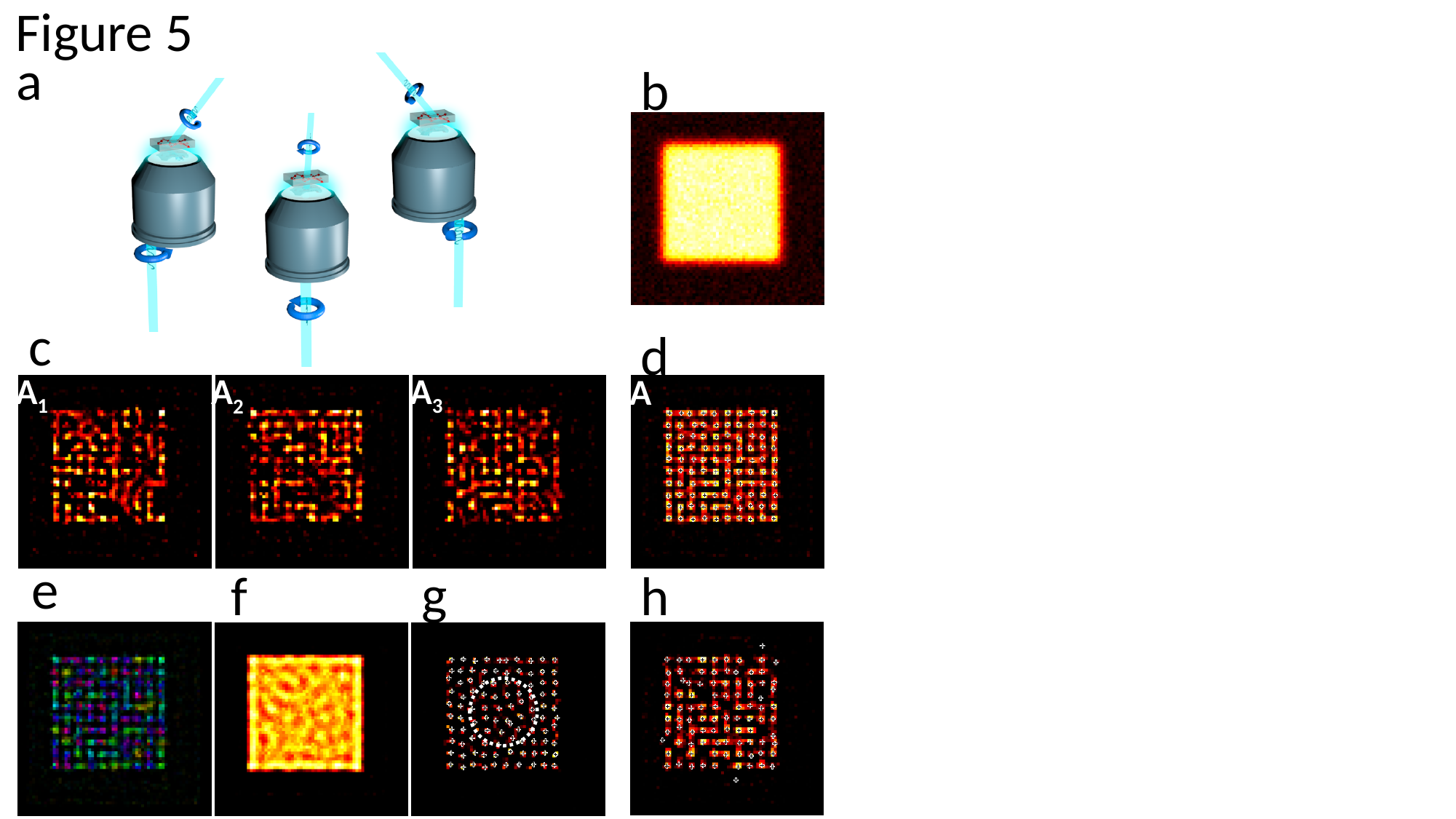

Figure 5
a
b
c
d
A_3_
A_1_
A_2_
A
e
g
f
h

### Slide 6
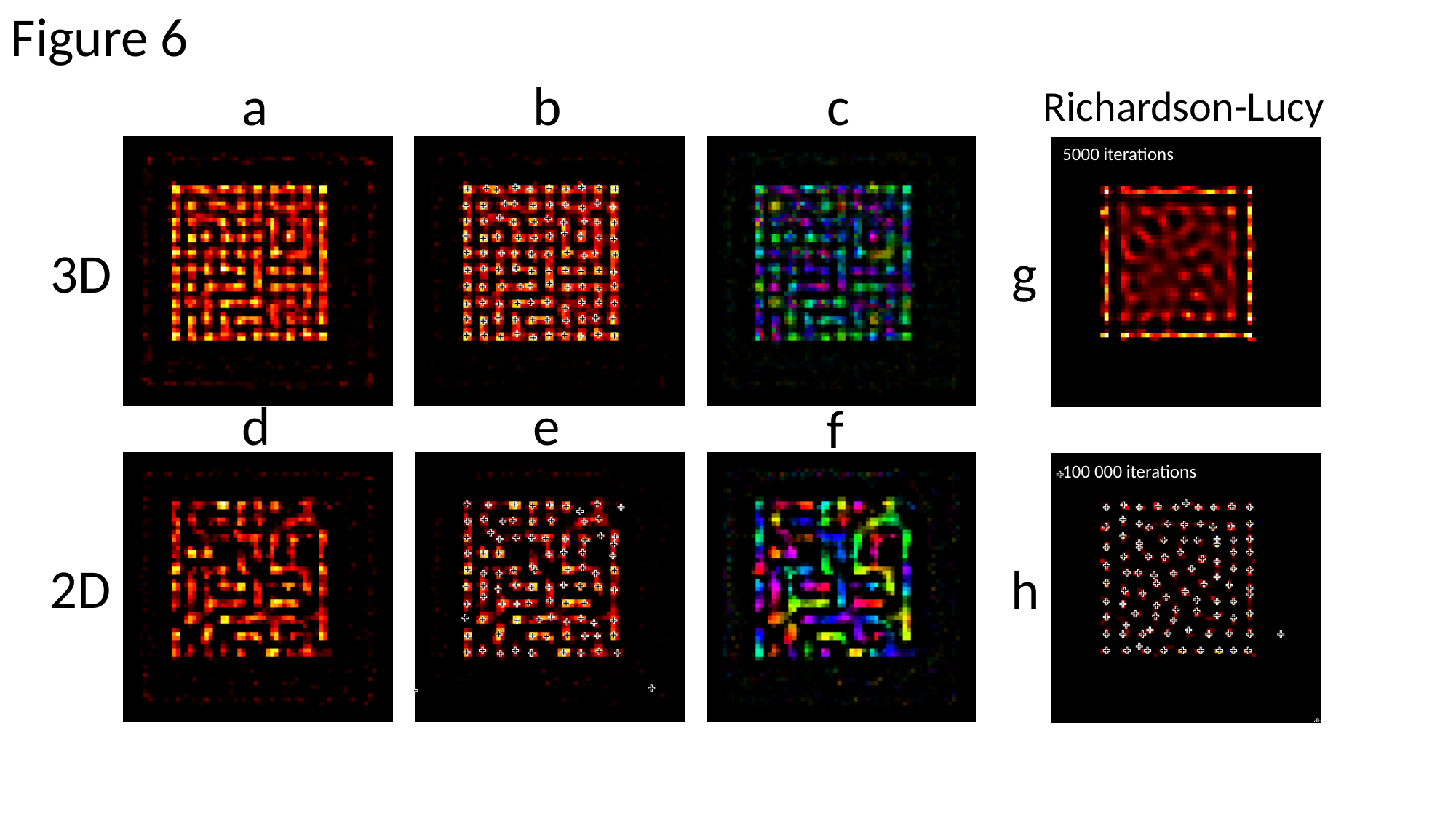

Figure 6
a
b
c
Richardson-Lucy
5000 iterations
3D
g
d
e
f
100 000 iterations
2D
h

### Slide 7
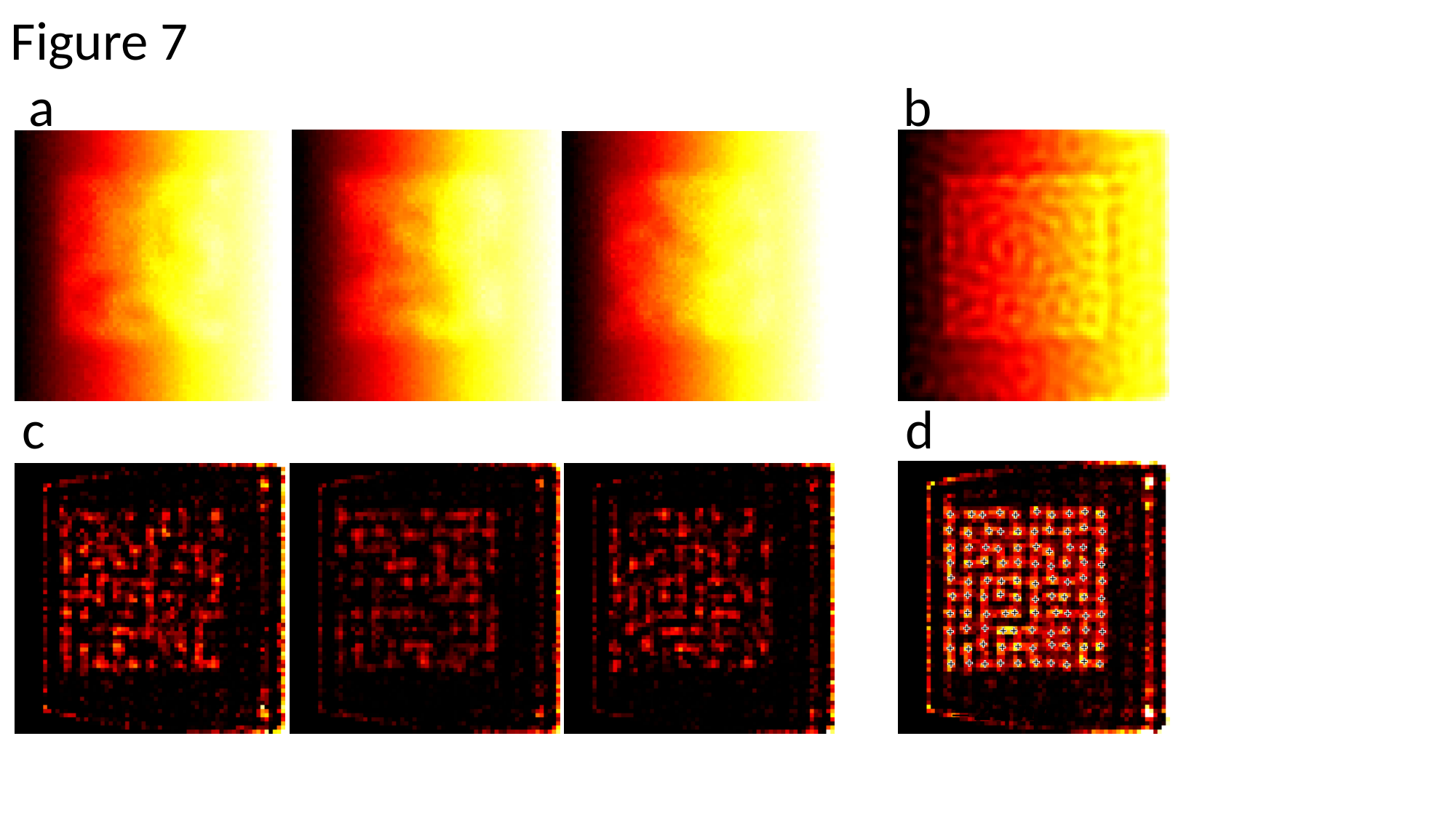

Figure 7
a
b
c
d

### Slide 8
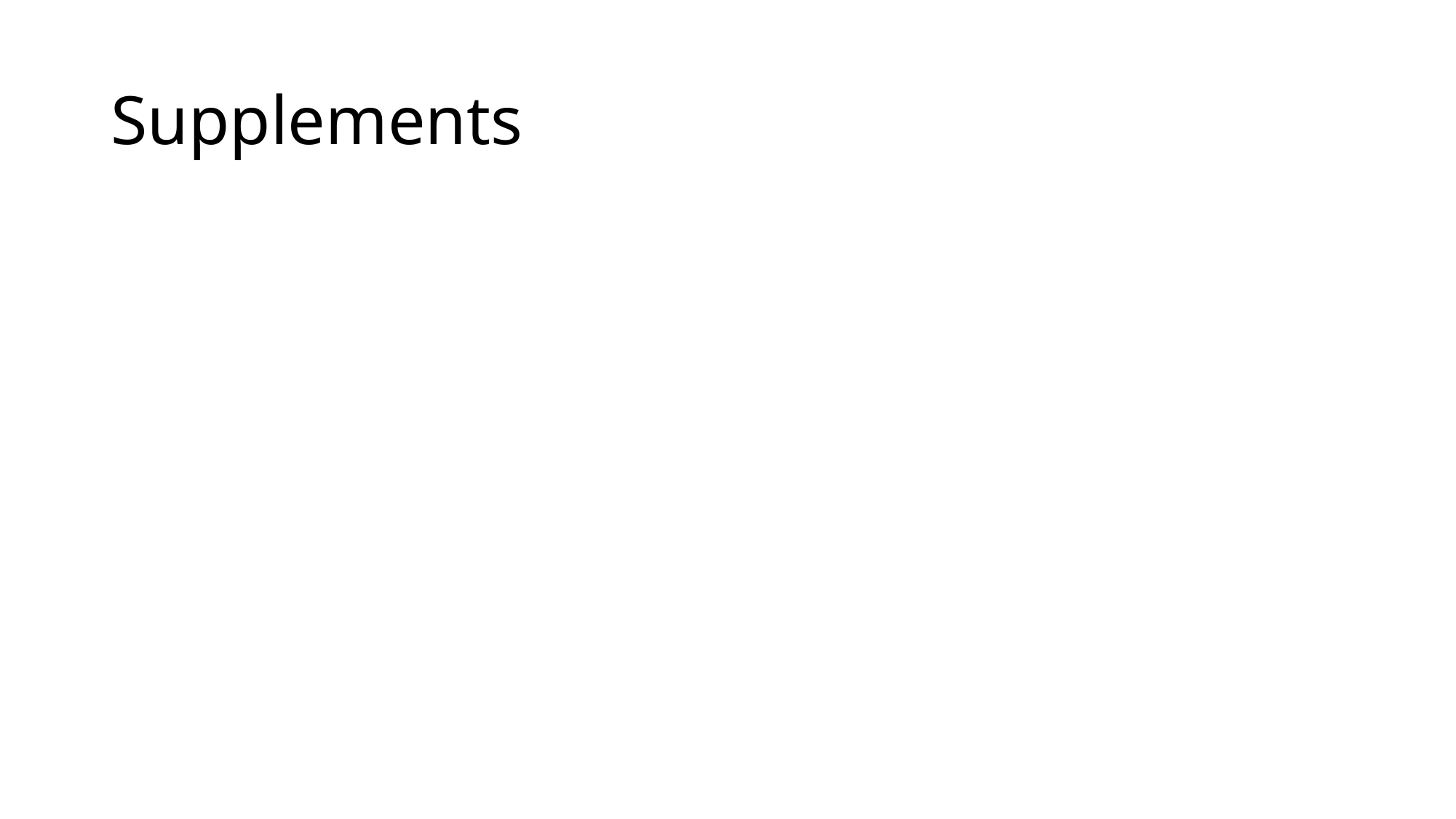

#### Supplements

### Slide 9
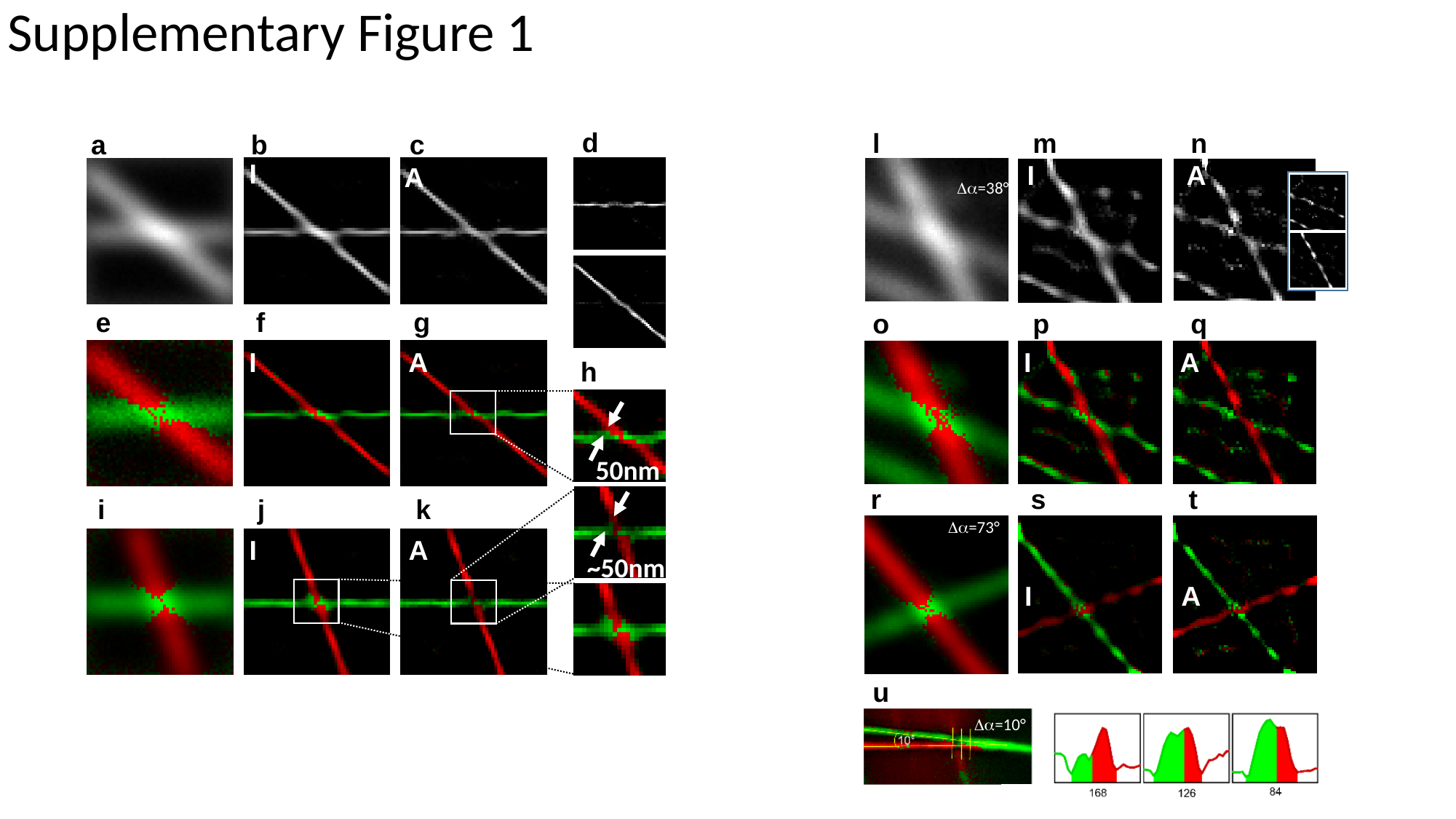

Supplementary Figure 1
d
l
m
n
a
b
c
c
I
I
A
A
Δα=38°
e
f
g
o
p
q
I
A
I
A
h
50nm
r
s
t
i
j
k
Δα=73°
I
A
~50nm
I
A
u
Δα=10°

### Slide 10
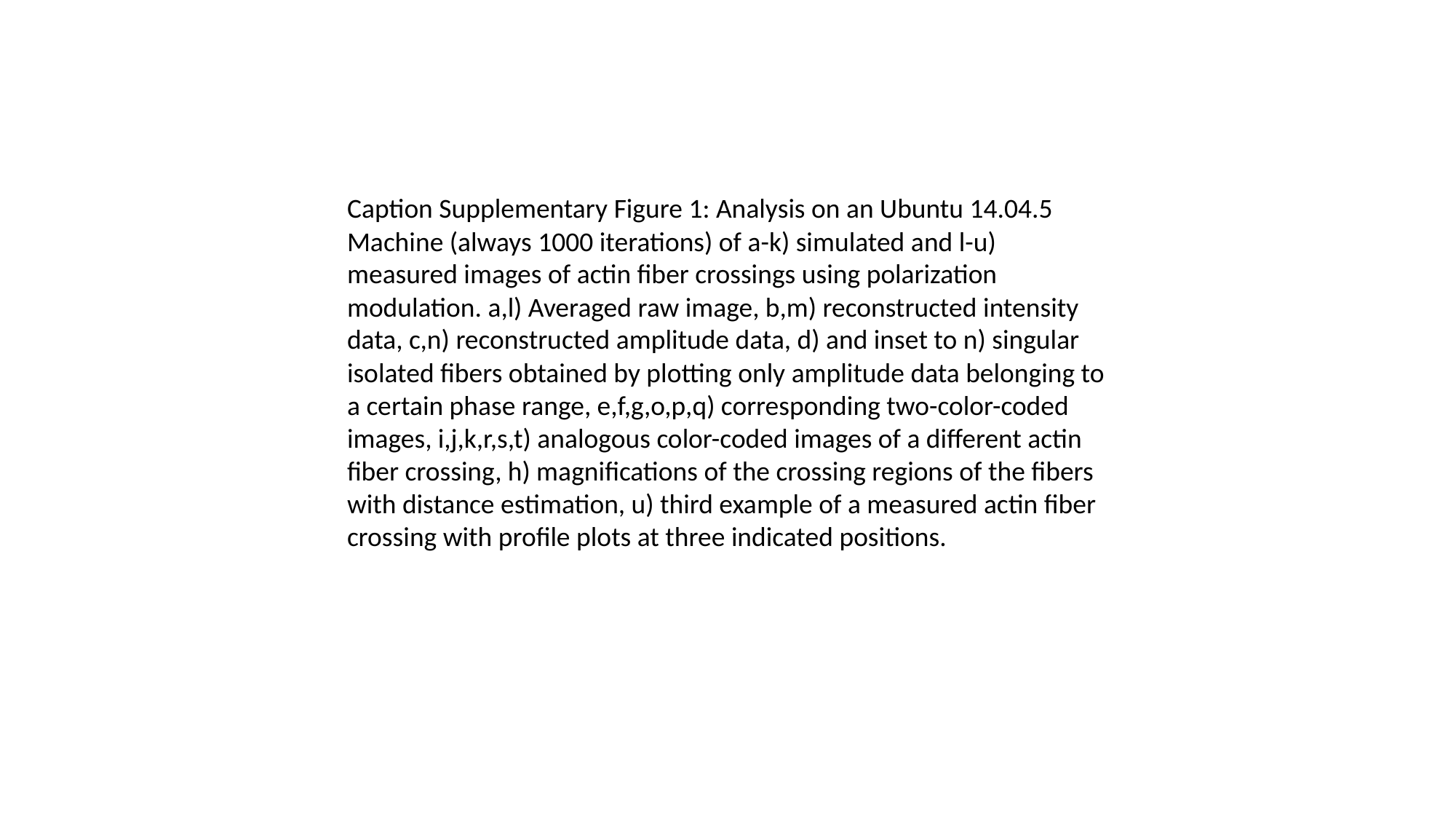

Caption Supplementary Figure 1: Analysis on an Ubuntu 14.04.5 Machine (always 1000 iterations) of a-k) simulated and l-u) measured images of actin fiber crossings using polarization modulation. a,l) Averaged raw image, b,m) reconstructed intensity data, c,n) reconstructed amplitude data, d) and inset to n) singular isolated fibers obtained by plotting only amplitude data belonging to a certain phase range, e,f,g,o,p,q) corresponding two-color-coded images, i,j,k,r,s,t) analogous color-coded images of a different actin fiber crossing, h) magnifications of the crossing regions of the fibers with distance estimation, u) third example of a measured actin fiber crossing with profile plots at three indicated positions.

### Slide 11
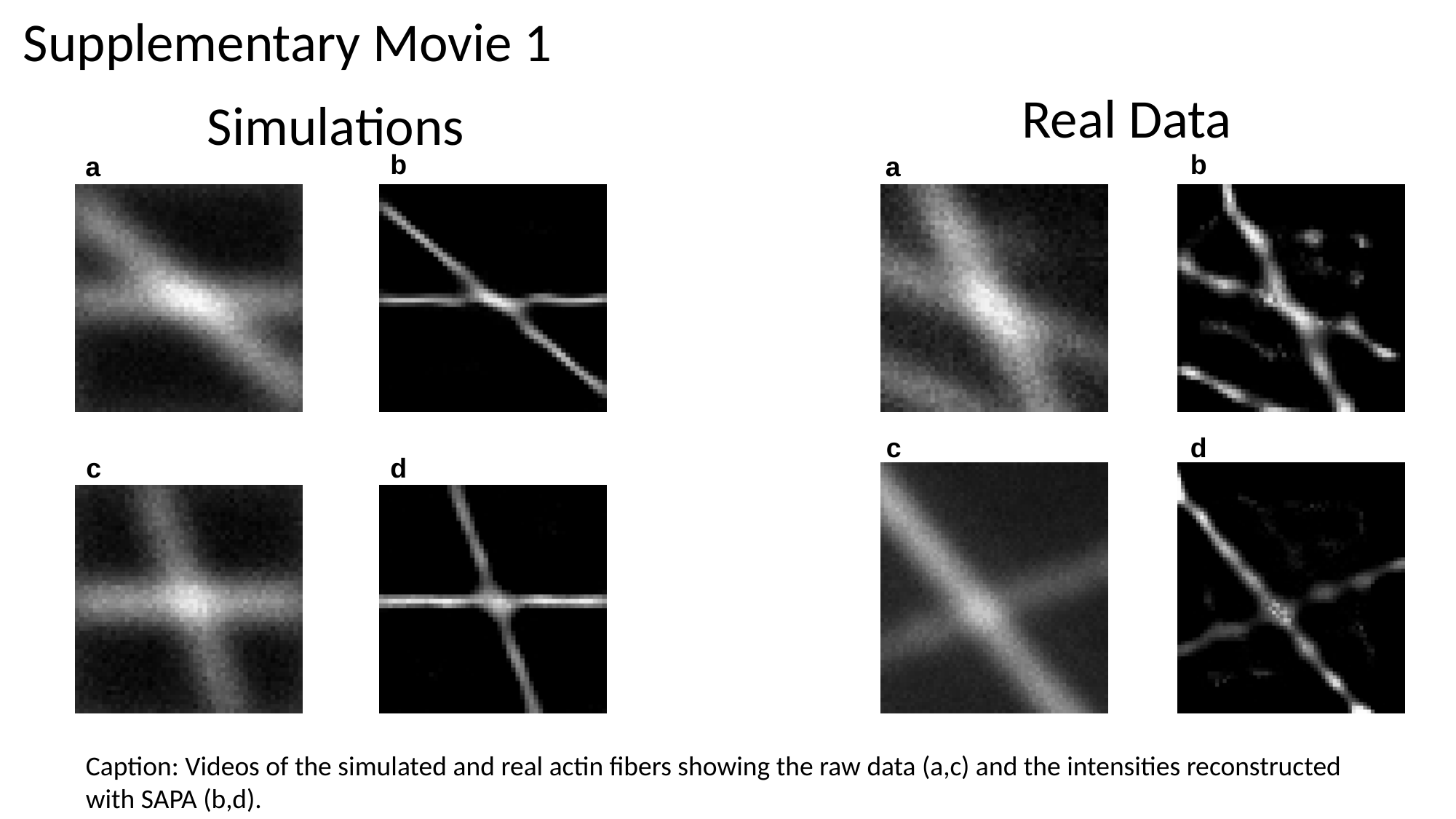

Supplementary Movie 1
Real Data
Simulations
b
b
a
a
c
d
c
d
Caption: Videos of the simulated and real actin fibers showing the raw data (a,c) and the intensities reconstructed with SAPA (b,d).

### Slide 12
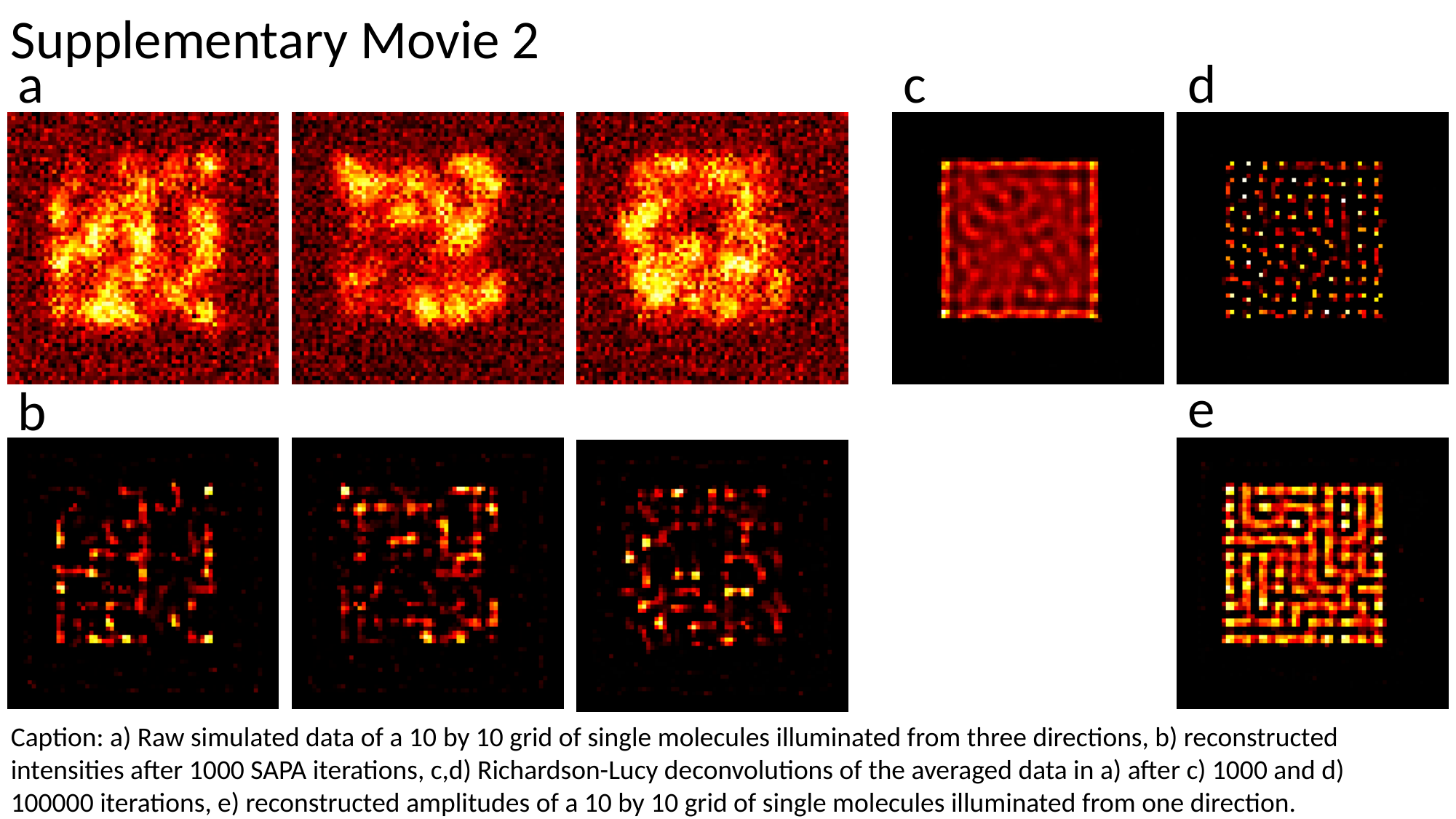

Supplementary Movie 2
d
a
c
e
b
Caption: a) Raw simulated data of a 10 by 10 grid of single molecules illuminated from three directions, b) reconstructed intensities after 1000 SAPA iterations, c,d) Richardson-Lucy deconvolutions of the averaged data in a) after c) 1000 and d) 100000 iterations, e) reconstructed amplitudes of a 10 by 10 grid of single molecules illuminated from one direction.

### Slide 13
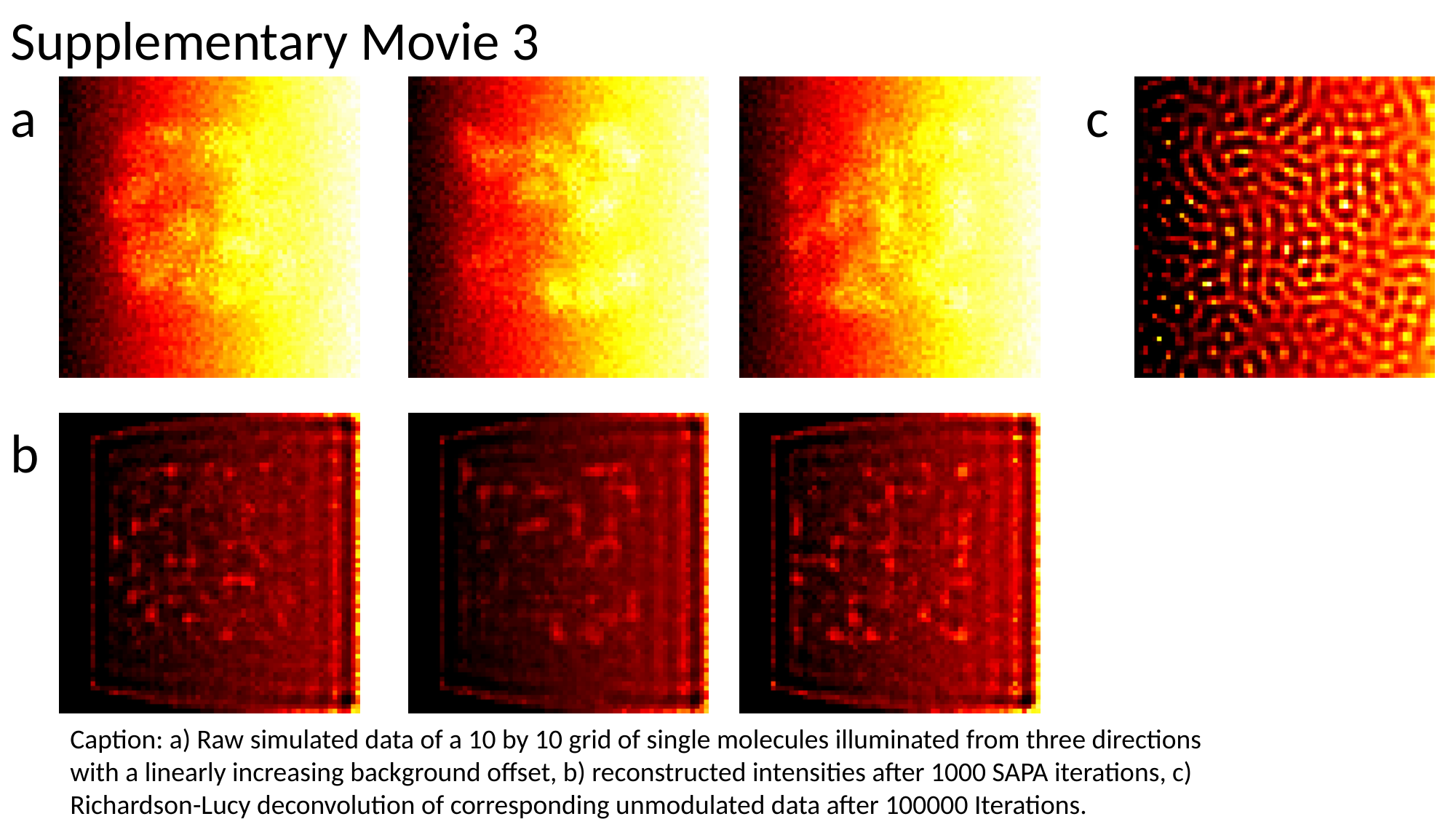

Supplementary Movie 3
c
a
b
Caption: a) Raw simulated data of a 10 by 10 grid of single molecules illuminated from three directions with a linearly increasing background offset, b) reconstructed intensities after 1000 SAPA iterations, c) Richardson-Lucy deconvolution of corresponding unmodulated data after 100000 Iterations.
